# Recent adverse mortality trends in Scotland: comparison with other high-income countries

**DOI:** 10.1101/542449

**Authors:** Lynda Fenton, Jon Minton, Julie Ramsay, Maria Kaye-Bardgett, Colin Fischbacher, Grant MA Wyper, Gerry McCartney

## Abstract

**Objective:** Gains in life expectancy have faltered in several high-income countries in recent years. We aim to compare life expectancy trends in Scotland to those seen internationally, and to assess the timing of any recent changes in mortality trends for Scotland.

**Setting:** Austria, Croatia, Czech Republic, Denmark, England & Wales, Estonia, France, Germany, Hungary, Iceland, Israel, Japan, Korea, Latvia, Lithuania, Netherlands, Northern Ireland, Poland, Scotland, Slovakia, Spain, Sweden, Switzerland, USA.

**Methods:** We used life expectancy data from the Human Mortality Database (HMD) to calculate the mean annual life expectancy change for 24 high-income countries over five-year periods from 1992 to 2016, and the change for Scotland for five-year periods from 1857 to 2016. One- and two-break segmented regression models were applied to mortality data from National Records of Scotland (NRS) to identify turning points in age-standardised mortality trends between 1990 and 2018.

**Results:** In 2012-2016 life expectancies in Scotland increased by 2.5 weeks/year for females and 4.5 weeks/year for males, the smallest gains of any period since the early 1970s. The improvements in life expectancy in 2012-2016 were smallest among females (<2.0 weeks/year) in Northern Ireland, Iceland, England & Wales and the USA and among males (<5.0 weeks/year) in Iceland, USA, England & Wales and Scotland. Japan, Korea, and countries of Eastern Europe have seen substantial gains in the same period. The best estimate of when mortality rates changed to a slower rate of improvement in Scotland was the year to 2012 Q4 for males and the year to 2014 Q2 for females.

**Conclusion:** Life expectancy improvement has stalled across many, but not all, high income countries. The recent change in the mortality trend in Scotland occurred within the period 2012-2014. Further research is required to understand these trends, but governments must also take timely action on plausible contributors.

**Strengths and limitations of this study:**

- The use of five-year time periods for comparison of life expectancy changes reduces the influence of year-to-year variation on observations.
- Examining long-term trends addresses concerns that recent life expectancy stalling may be over-emphasised due to notably large gains in the immediately preceding period.
- The international comparison was limited to the 24 high-income countries for which data were readily available for the relevant period.
- Analysis of trend data will always be sensitive to the period selected, however segmented regression of the full period of mortality rates available offers an objective method of identifying the timing of a change in trend.

## Background

Mortality rates have steadily declined, and life expectancy has improved, in most high-income countries since 1945.^1,2^ There have been previous exceptions to this general trend, including the countries of Eastern Europe during the 1990s.^1,3^ Recently there have been a series of reports suggesting that mortality improvements are now faltering, or even reversing, for the USA, the UK, and much of continental Europe, since around 2011.^4–6^

Contextualising current mortality trends within those that have been observed previously and internationally can support a proportionate public health response, and identify comparator countries or periods to assist future investigation of causal hypotheses. International comparison of changes in life expectancy across a single year (2014 to 2015) found that life expectancy declined in 8 out of 18 high-income countries, including the UK.^4^ However, the short-run trends in mortality data, even at national level, can vary substantially from year-to-year and observations may be therefore by sensitive to the comparison period.^7^ Comparison of the most recent six years to the preceding six years found that, of 20 countries, the UK had had the largest life expectancy slow-down for females, and the second largest for males.^5^ This however, does not allow identification of which period was exceptional: the previous gains or the current slow-down.

Among the UK countries Scotland has the lowest life expectancy, with a period life expectancy at birth in 2015-2017 which was 2.0 years lower for women, and 2.5 years lower for men than that observed in England.^8^ Analysis by the UK Office for National Statistics (ONS) found that a slowdown in mortality rates has been seen in all four UK countries in 2011-2016 compared to 2006-2011, but that Scotland experienced the least stalling for women, and second least after Northern Ireland for men.^6^

Several hypotheses have been proposed to explain recent changes in life expectancy trends. Cohort effects, whereby a particular generation is at a higher risk of mortality, may be important if that generation is now reaching an age where it contributes more to overall mortality and life expectancy.^2,9^ Other possibilities are that there is an interaction between period effects (such as policy changes or infectious disease epidemics) and vulnerabilities within a cohort such that mortality for that group increases. This has been observed for specific causes of death in Scotland and the USA (suicide, drug-related deaths and alcohol).^10–13^

There has been an apparent polarisation of the debate regarding causes of recent adverse mortality trends, between explanations emphasising influenza, and those concerned with the impacts of austerity.^14–18^ It may be that this split is in part attributable to studies seeking the answers to different questions (for example the causes of high numbers of deaths in short periods of time versus stalling of overall life expectancy over longer periods) and in variable comparator, or baseline, periods employed. Causal investigation would be strengthened by clear description of the nature, scale and timing of the phenomenon we are seeking to explain.

This study aims to describe the nature, scale and timing of changes in mortality in Scotland, and to compare these to those seen internationally, as an early step in understanding their causes.

## Methods

We report our results in accordance with the RECORD guideline.^19^

### Data

We used population data from the Human Mortality Database (HMD)^20^ for life expectancy analyses. Segmented regression analysis of age-standardised mortality rates used data held by National Records of Scotland (NRS). All analyses were undertaken for males and females separately.

### Life expectancy: average annual change in five-year periods

Period life expectancy figures for Scotland for each single year between 1855 and 2016 were extracted. For international comparisons, data were obtained for all high-income countries within the HMD which provided data for 2016 at the time of extraction^1^. The mean annual change in life expectancy (in weeks) for five-year periods running back from 2016 was calculated for each country. A sensitivity analysis using rolling five-year time periods rather than set periods from 2016 backwards was also undertaken.

### Age-standardised mortality rates: segmented regression

We calculated directly age-standardised mortality rates per 100,000 population for rolling four-quarter periods for Scotland using the 2013 European Standard Population for the entire time period (Q1 1990 to Q2 2018). Population estimates were calculated for each four-quarter period by interpolating the mid-year estimates. Data points are labelled by their final quarter, so quarter 1 (Q1) 2016 represents the mortality rate for 2015 Q2, Q3 and Q4 combined with 2016 Q1. Quarterly-rolling rates were used in order to increase the number of data points available to the model. In order to identify the point in the time series at which a change in trend occurred, we undertook segmented regression in R using the ‘segmented’ package. ^21,22^ We used the Davies test for the existence and statistical significance of a breakpoint. We used the segmented test, which treats the whole time series as continuous, to identify the breakpoint and standard error. The results of the segmented test were interpreted as identifying the quarterly data point within which the breakpoint fell. In this way a result of 2014.374 falls within quarter 2 of 2014, and the data which correspond to this quarter represent the ‘year’ quarter 3 2013 to quarter 2 2014, hence the year to 2014 Q2 is interpreted as the best estimate of when a change in trend occurred. Ninety-five percent confidence intervals for the breakpoint were calculated from the standard error of this estimate. We used the segmented test to examine one and two break point models and compared model fit using Akaike Information Criterion (AIC) and Bayesian Information Criterion (BIC) values. Segmented regression models were produced separately for all males, all females and for males and females divided into under 75 year and 75+ year age groups.

## Results

### Life expectancy trends

Period life expectancy at birth for men and women in Scotland increased from 44 years for women and 41 years for men in 1855 to 81 years for women and 77 years for men in 2016, based on single-year estimates. Throughout this period women had longer life expectancies than men. The trend up to around 1945 was substantially more unstable than in later years, but there was a general improvement, especially after 1890. From 1950 the degree of year-to-year variability reduced and there was a slower, steady improvement.

The mean annual change in life expectancy observed in Scotland in five-year periods (1857 and 2016) shows that the largest gains were made in the periods following declines in life expectancy (e.g. 1942-1946) (Figure 1). From 1997-2011 each period saw steady gains for females (range 9.8-11.0 weeks/year) and males (range 14.1-17.3 week/year). In the period 2012-2016, only small mean life expectancy improvements were observed: 2.5 weeks/year for females and 4.5 weeks/year for males. This represents the smallest average annual increase for women since 1937-41, and for men since 1972-76. A sensitivity analysis (Appendix figure 1) using rolling five-year periods identifies similar periods of slow life expectancy gain, showing results are not dependent on the selection of particular start and finish years.

**Figure 1:**
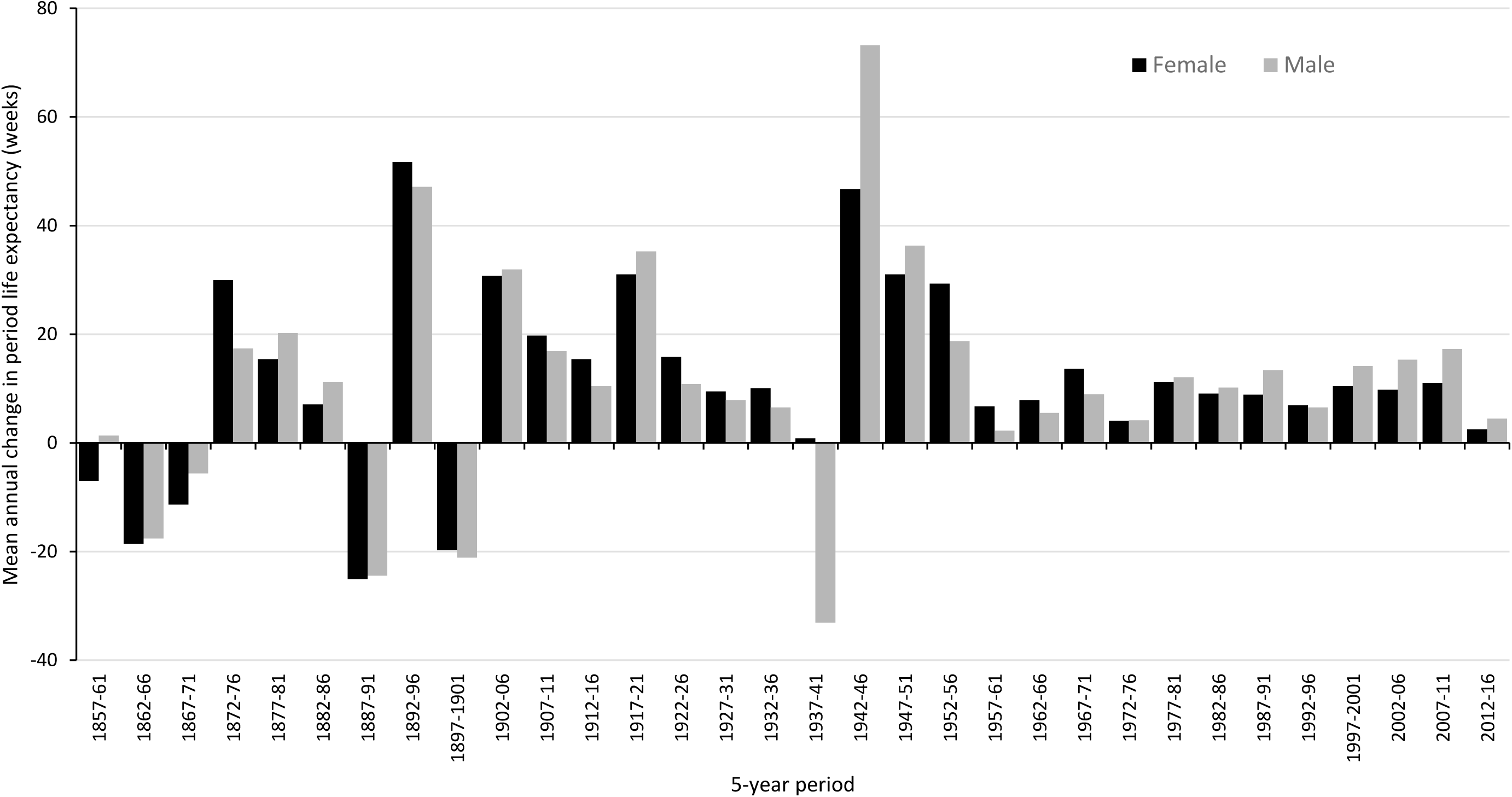
Mean annual change in period life expectancy at birth (weeks) for five-year periods, men and women, Scotland (civilian population), 1857-2016.

To identify the nations and time periods with the greatest change in life expectancy trends over the last three decades, the mean annual changes in life expectancy (in weeks) for all 24 high-income countries with HMD data available to 2016 are shown in Figures 2 and 3, for females and males respectively. Nearly all countries saw mean increases in life expectancy across all five-year time periods, with the exceptions among females being: Northern Ireland (2012-2016), Iceland (1997-2001) Latvia (1992-1996), and Lithuania (2002-2006), and among males: Iceland and USA (2012-2016), Latvia (1992-1996) and Lithuania (1992-1996 and 2002-2006).

**Figure 2:**
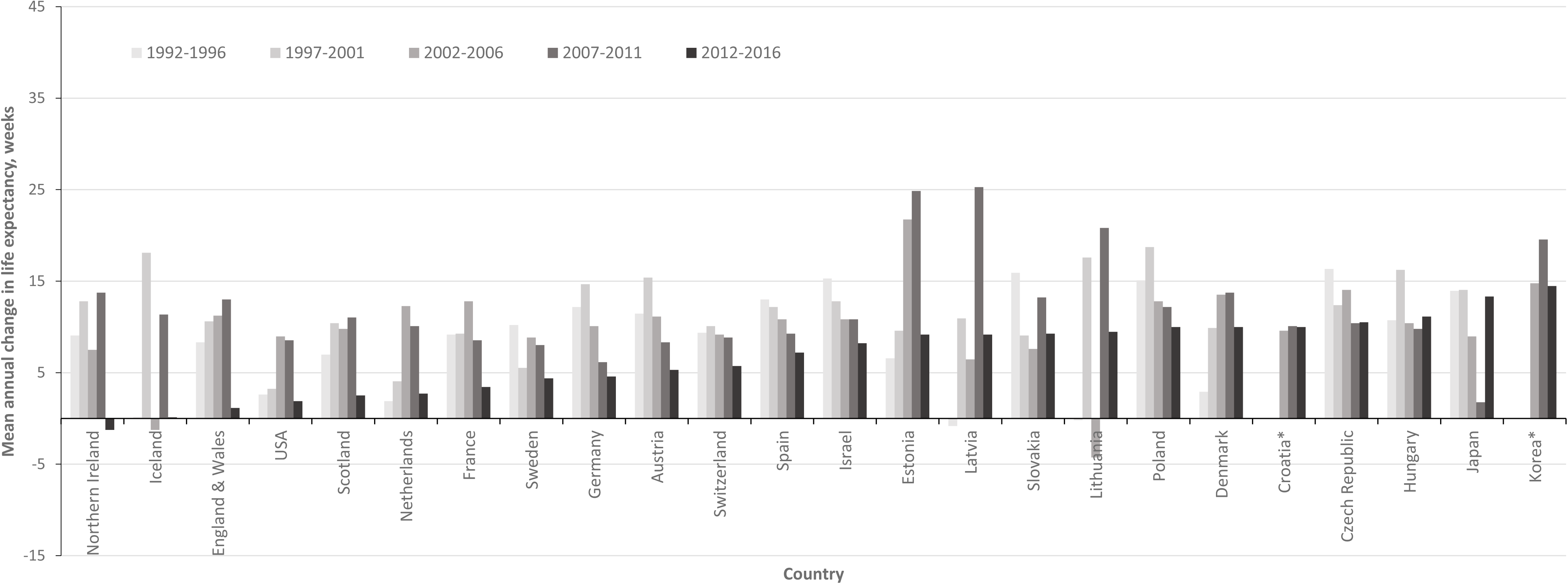
Mean annual change in female life expectancy at birth (weeks), for five-year periods 1991-2016, by country. *no data available for Croatia and Korea for periods prior to 2002.

**Figure 3:**
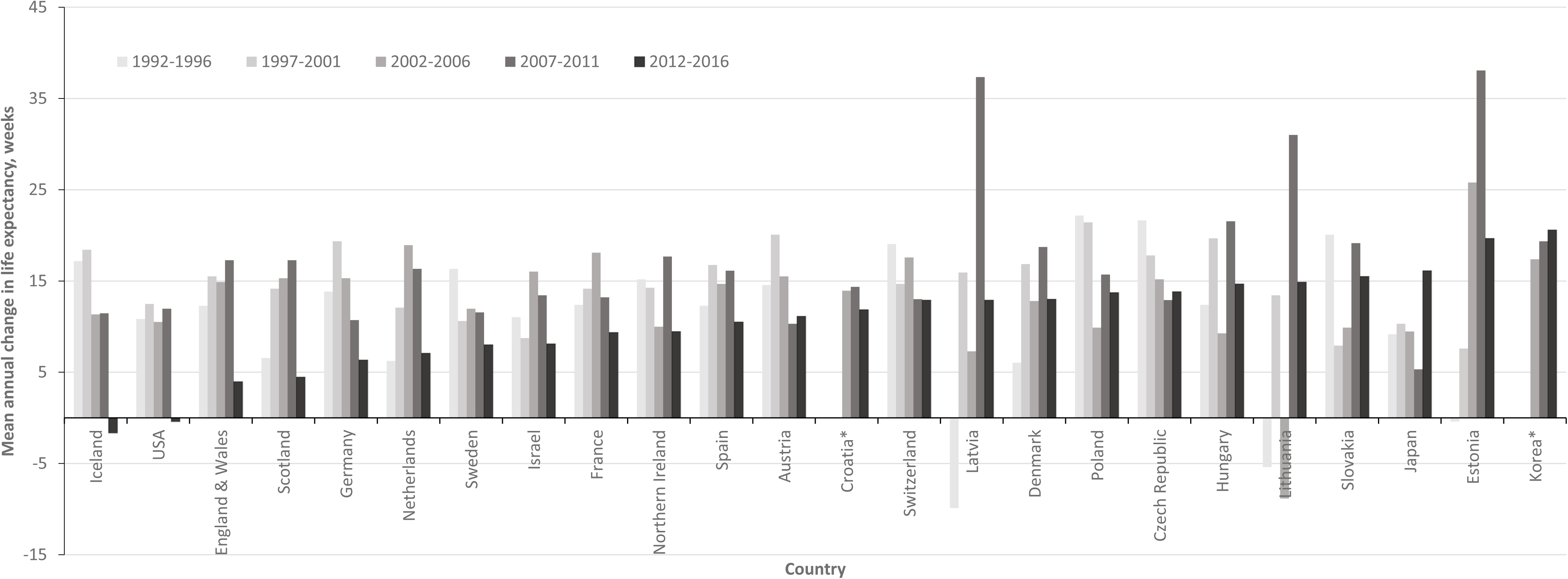
Mean annual change in male life expectancy at birth (weeks), for five-year periods 1991-2016, by country. *no data available for Croatia and Korea for periods prior to 2002.

For females, the range of mean life expectancy change in 2012-2016 was −1.3 to 14.5 weeks/year (interquartile range 3.3 to 10.0 weeks/year). Nine countries saw mean gains of less than five weeks/year: Northern Ireland (−1.2 weeks/year), Iceland (0.1 weeks/year), England & Wales (1.1 weeks/year), USA (1.9 weeks/year), Scotland (2.5 weeks/year), the Netherlands (2.7 weeks/year), France (3.4 weeks/year) and Sweden (4.4 weeks/year), and Germany (4.6 weeks/year). Seven countries had mean gains of 10 weeks per year or more: Poland (10.0 weeks/year), Denmark (10.0 weeks/year), Croatia (10.0 weeks/year), Czech Republic (10.5 weeks/year), Hungary (11.1 weeks/year), Japan (13.3 weeks/year) and Korea (14.5 weeks/year). Life expectancy gain was smaller in 2012-2016 than the preceding 5 years for all countries except the Czech Republic, Hungary and Japan (Figure 2).

Amongst males, the range of mean life expectancy change in 2012-2016 was −1.7 to 20.6 weeks/year (interquartile range 7.8 to 14.0 weeks/year). Four countries had mean gains of less than five weeks/year: Iceland (−1.7 weeks/year), USA (−0.4 weeks/year), England & Wales (4.0 weeks/year), and Scotland (4.5 weeks/year). Fourteen countries had gains of 10 weeks/year or more: Spain (10.5 weeks/year), Austria (11.1 weeks/year), Croatia (11.9 weeks/year), Switzerland (12.9 weeks/year) Latvia (12.9 weeks/year), Denmark (13.0 weeks/year), Poland (13.7 weeks/year), Czech Republic (13.8 weeks/year), Hungary (14.7 weeks/year), Lithuania (14.9 weeks/year), Slovakia (15.5 weeks/year), Japan (16.1 weeks/year), Estonia (19.7 weeks/year) and Korea (20.6 weeks/year). The increases for the 2012-2016 were smaller than in 2007-2011 for all countries except Japan and Korea (Figure 3).

### Segmented regression

Figure 4 shows the rolling four-quarter age standardised mortality rates (ASMRs), by sex, for Scotland for all ages. Over the period (1990 Q1 – 2018 Q2), the ASMR per 100,000 population fell from 2,114 to 1,355 for males, and from 1,386 to 1,025 for females. Males had a higher mortality rate than females throughout the series, although this gap narrowed over time. The steadiest period of decline in mortality rates appeared to be from 2004 to around 2011, with the periods before and after this showing variation between slow improvements, worsening of mortality rates, and faster improvements. The mortality rates for those aged 75+ years showed greater variability than those in the younger age group.

**Figure 4:**
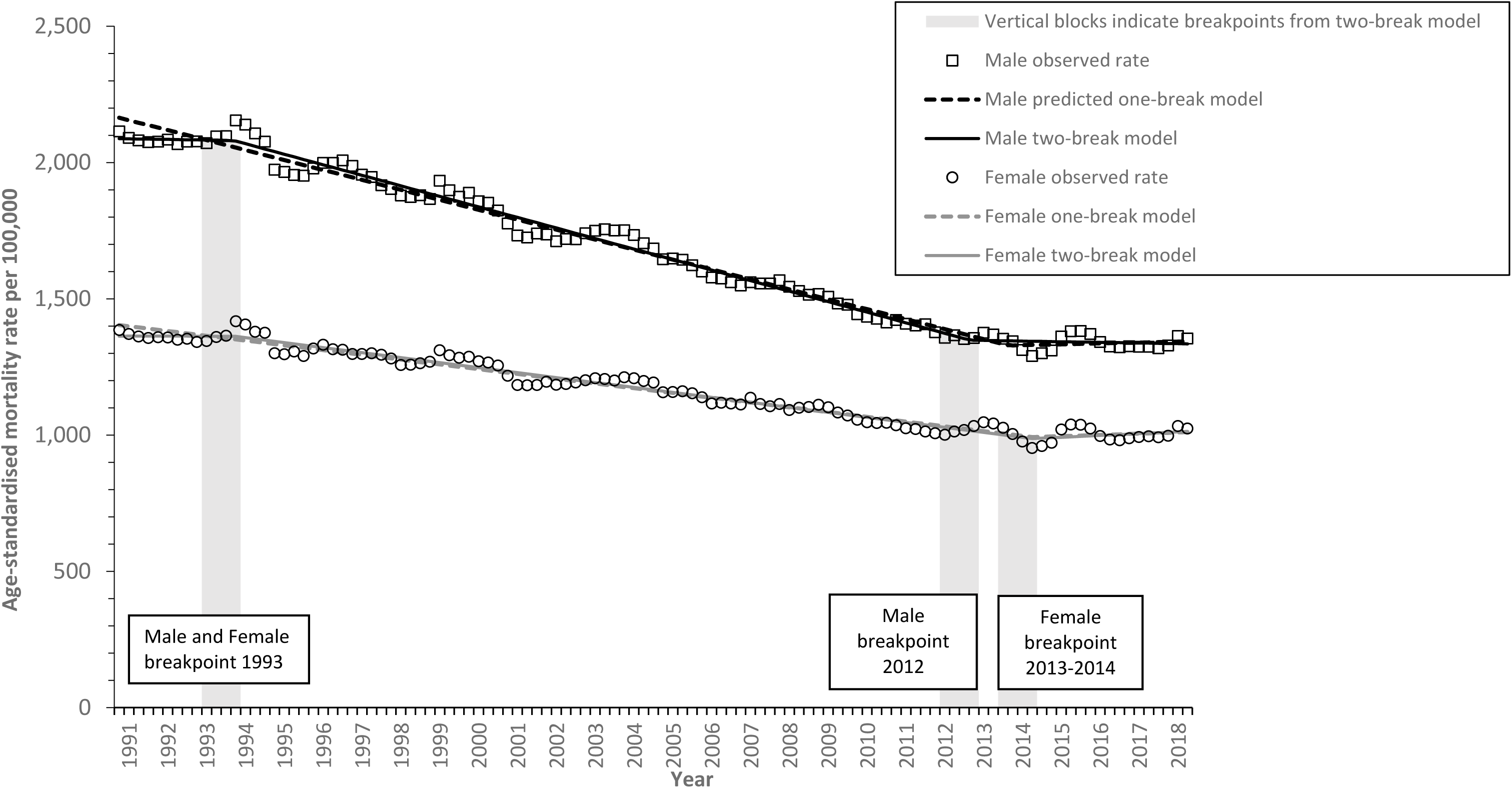
Age-standardised rolling four-quarterly mortality rates, with segmented regression models fitted, Scotland, 1990-2018.

The Davies test for the existence of a change in the slope identified a statistically significant change (p<0.01) for males and females, and both age groups tested. For all groups the breakpoint identified by the Davies test fell within the period 2012-2014 (see Table 1). The segmented model provides a more precise approach to estimation of the timing of the breakpoint. The date estimates from the one-break segmented model corresponded to those identified by the Davies test for all groups, to within 0.2 years. One and two-break models were run for all groups; both AIC and BIC were lower for the two-break models, indicating that these are a better fit, hence the results below report the two-break model findings.

**Table 1:**
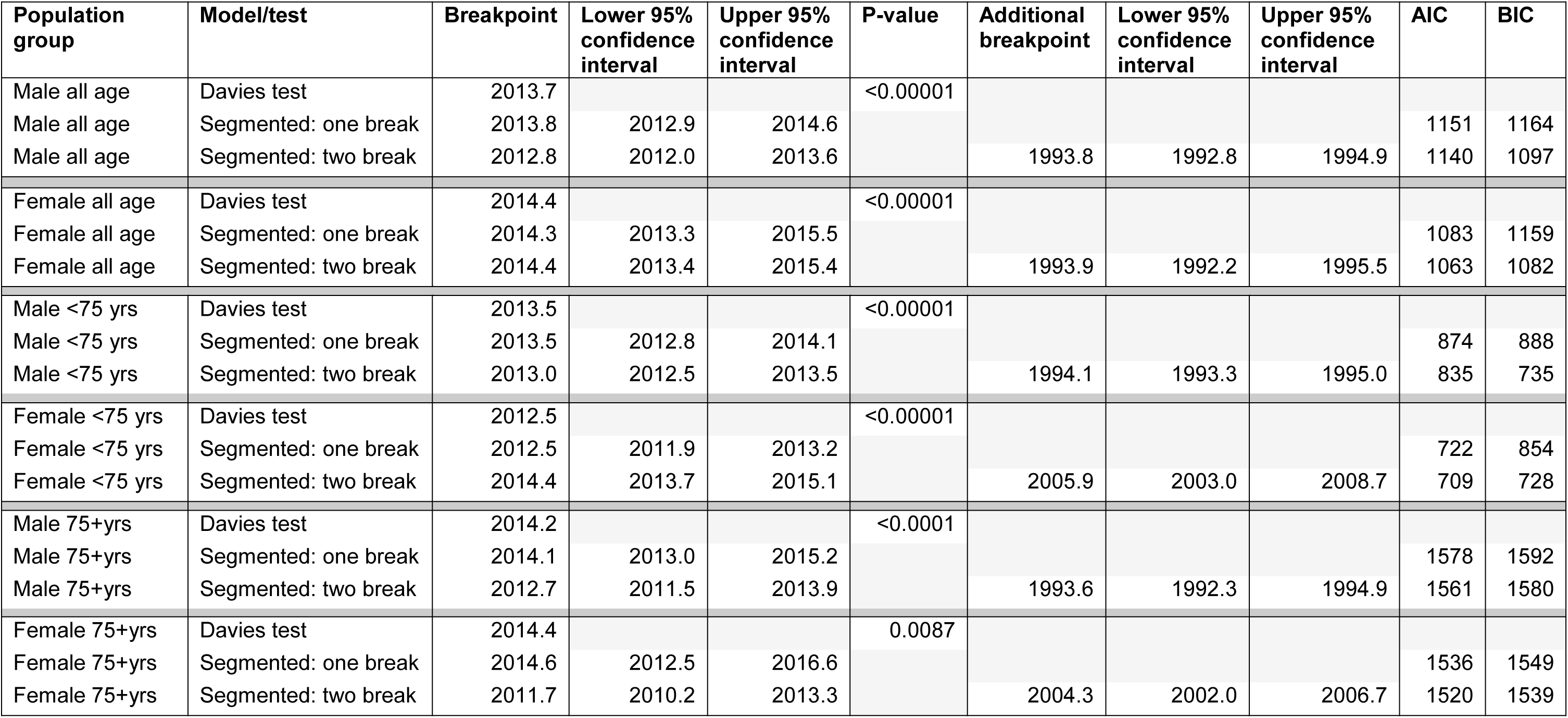
Summary of results of segmented regression by population group and model/test.

| Population group | Model/test | Breakpoint | Lower 95% confidence interval | Upper 95% confidence interval | P-value | Additional breakpoint | Lower 95% confidence interval | Upper 95% confidence interval | AIC | BIC |
| --- | --- | --- | --- | --- | --- | --- | --- | --- | --- | --- |
| Male all age | Davies test | 2013.7 |  |  | <0.00001 |  |  |  |  |  |
| Male all age | Segmented: one break | 2013.8 | 2012.9 | 2014.6 |  |  |  |  | 1151 | 1164 |
| Male all age | Segmented: two break | 2012.8 | 2012.0 | 2013.6 |  | 1993.8 | 1992.8 | 1994.9 | 1140 | 1097 |
| Female all age | Davies test | 2014.4 |  |  | <0.00001 |  |  |  |  |  |
| Female all age | Segmented: one break | 2014.3 | 2013.3 | 2015.5 |  |  |  |  | 1083 | 1159 |
| Female all age | Segmented: two break | 2014.4 | 2013.4 | 2015.4 |  | 1993.9 | 1992.2 | 1995.5 | 1063 | 1082 |
| Male <75 yrs | Davies test | 2013.5 |  |  | <0.00001 |  |  |  |  |  |
| Male <75 yrs | Segmented: one break | 2013.5 | 2012.8 | 2014.1 |  |  |  |  | 874 | 888 |
| Male <75 yrs | Segmented: two break | 2013.0 | 2012.5 | 2013.5 |  | 1994.1 | 1993.3 | 1995.0 | 835 | 735 |
| Female <75 yrs | Davies test | 2012.5 |  |  | <0.00001 |  |  |  |  |  |
| Female <75 yrs | Segmented: one break | 2012.5 | 2011.9 | 2013.2 |  |  |  |  | 722 | 854 |
| Female <75 yrs | Segmented: two break | 2014.4 | 2013.7 | 2015.1 |  | 2005.9 | 2003.0 | 2008.7 | 709 | 728 |
| Male 75+yrs | Davies test | 2014.2 |  |  | <0.0001 |  |  |  |  |  |
| Male 75+yrs | Segmented: one break | 2014.1 | 2013.0 | 2015.2 |  |  |  |  | 1578 | 1592 |
| Male 75+yrs | Segmented: two break | 2012.7 | 2011.5 | 2013.9 |  | 1993.6 | 1992.3 | 1994.9 | 1561 | 1580 |
| Female 75+yrs | Davies test | 2014.4 |  |  | 0.0087 |  |  |  |  |  |
| Female 75+yrs | Segmented: one break | 2014.6 | 2012.5 | 2016.6 |  |  |  |  | 1536 | 1549 |
| Female 75+yrs | Segmented: two break | 2011.7 | 2010.2 | 2013.3 |  | 2004.3 | 2002.0 | 2006.7 | 1520 | 1539 |

The two-break model for all ages identified the first breakpoint in the year to 1993 Q4 for both males (95% confidence interval (CI): year to 1992 Q4 - year to 1994 Q4) and females (95% CI: year to 1992 Q1 – year to 1995 Q2). A second breakpoint for males was identified in the year to 2012 Q4 (95% CI: year to 2012 Q1 – year to 2013 Q3), and for females in the year to 2014 Q2 (95% CI: year to 2013 Q2 – year to 2015 Q2). The change in trend indicated by these breakpoints is shown in Figure 4; the break in 1993 indicates a change from a period of slower mortality improvement to a period of faster improvement and the later breaks in 2012 and 2014 (males and females respectively) indicate a change to much slower gains.

Among those aged under 75 years, the results of the two-break model suggested that the later change in trend occurred approximately 18 months earlier in males (year to 2012 Q4) than in females (year to 2014 Q2), with the 95% confidence intervals for the estimates not overlapping (see Table 1). For those aged 75+ years the estimate for males (year to 2012 Q3) was one year later than for females (year to 2011 Q3), but the 95% confidence intervals for the estimates overlap.

Among males, the estimate of the later breakpoint of the two-break model was similar for those aged under 75 years and 75+ years (year to 2012 Q4 for both groups). For females the later breakpoint occurred nearly 3 years later in those aged under 75 years (year to 2014 Q2) than those aged 75+ years (year to 2011 Q3), with the 95% confidence intervals not overlapping.

## Discussion

### Principal findings

The increase in life expectancy in Scotland since 1855 has occurred at different rates over time. Over the first one hundred years examined here, there were periods of rapid increase, but also notable declines. Since 1957, however, there has been a pattern of smaller, steadier increases in life expectancy for both males and females. The life expectancy gains between 2012 and 2016 are amongst the smallest seen in this later period, with average increases of only 2.5 weeks/year for women and 4.5 weeks/year for men.

Of the 24 high-income countries for which data were available, nearly all had smaller life expectancy gains in 2012-2016 than in the immediately preceding period. Japan and Korea are notable exceptions, and in Japan there was a substantial slow-down in life expectancy gains in the period 2007-2011, followed by a resumption of gains at the level previously seen. Among the countries with a stalling of life expectancy gains in 2012-2016 there is large variation in the mean change observed, and in the scale of the difference between periods. Iceland, the USA, England & Wales, Scotland, and, for females, Northern Ireland, had the smallest gains in 2012-2016 and the most marked stalling. In general, the countries of Western Europe saw smaller gains in 2012-2016, with some degree of slow-down, compared to countries of Eastern Europe, where steadier gains have been maintained. Denmark is notable for having maintained mean life expectancy gains of around 10 weeks/year among females across the period 1997-2016, and even greater gains among males.

The two-break segmented regression model suggests that mortality trends changed to a pattern of more rapid improvement for both males and females in the year to 1993 Q4. In the year to 2012 Q4 for males, and 2014 Q2 for females, the trend in mortality rates changes again, with an increase thereafter. For all the models and groups tested, a negative turning point in mortality rates was consistently identified within the period 2011 to 2015.

### Strengths and limitations

Using life expectancy and age-standardised mortality rates ensures that the analyses in this paper are not prone to confounding by changes in the age structure of the population. We used all-cause mortality rates, thus avoiding difficulties due to competing causes of death and coding uncertainties. We performed sensitivity analyses on the periodisation of the average gain in life expectancy comparisons to identify the potential for the findings to be affected by the selection of a particular start date for the analyses.

The use of single-year life expectancy estimates from the HMD allowed international comparison; it should be noted that these data differ slightly from life expectancy estimates published by NRS using 3-yearly rolling averages. The international analysis is limited to the range of countries for which data were available through the HMD at the time of extraction. We were only able to conduct segmented regression employing four-quarter rolling mortality rates for Scotland, as we did not have access to equivalent data for other countries. We acknowledge that the confidence intervals presented for segmented regression may underestimate the true uncertainty, as the nature of the rolling quarterly mortality rate estimates means that the data points aren’t discrete.

Whilst other studies have focused changes in mortality between single years (particularly 2014 to 2015), we were explicit in seeking to describe the longer-term mortality trends, and therefore employed five-year time periods for comparison to reduce the influence of year-to-year variation on observations. By extending life expectancy gain comparisons back over a longer time period we have sought to address concerns that the stalling of life expectancy in the most recent period may be over-emphasised due to notably large gains in the immediately preceding period.

### How this fits

Our overall findings are consistent with those of others, and the recent stalling of life expectancy gains across many high-income countries is now well recognised.^4–7^ Other analyses have emphasised the recent reduction in mortality improvements relative to those seen in the immediately preceding period.^5^ We have shown that relatively large life expectancy gains were seen for both males and females in Scotland in the preceding 15 years (1997-2011), but that even before this gains as small as those seen recently have not been observed since at least the early 1970s. Comparison of mortality trends within the UK suggests that the stalling seen in Scotland may not be as severe as that seen in England and Wales.^6^ Our findings confirm this, but allow us to place this difference within a wider international context which shows that the changes seen in Scotland are still more severe than those observed in many other high-income countries. The timing of a change in overall mortality trends found in this analysis is broadly consistent with that observed in England, where a breakpoint for females was found in the year to 2014 Q2, and the year to 2012 Q1 for males.^23^ Some differences are seen when data are age-stratified, with an earlier breakpoint observed in England for males <75 years and females 75+ years.

### Meaning – explanations and implications

Various hypotheses have been proposed to explain these trends, in particular the period effects of influenza and of economic austerity, and cohort effects, such as the impact on mortality risk of population cohorts with a high prevalence of obesity. It seems likely that factors common to all of the countries displaying similar trends, and absent in countries without the change in trend, are causal. It is also likely that several factors acting together are relevant to explaining the trends, whether that is some aspect of the context (such as the underlying political economy within a country) or two specific factors interacting. Many of the hypotheses proposed thus far are not mutually exclusive, but that does not mean that all the factors suggested are causal or have the same importance. It is possible that influenza and political economy explanations are both causal, with interactions between population vulnerability, social and health care pressures, and influenza.

The global financial crisis of 2008 led to a marked economic recession in many countries, and given that unemployment and income are important determinants of health,^24^ the potential for the crisis to adversely impact on mortality was highlighted early.^25^ However, the evidence around the impact of economic recession on health and mortality of populations, rather than individuals, is complex and contested.^26^ The response to this financial crisis, across many countries, was to implement a range of austerity policies whereby public spending was reduced in the pursuit of balanced budgets. As a result many public services experienced substantial reductions in their budgets and public sector wages and income transfers to lower income groups were frequently reduced in real terms. There is evidence that this impacted on a range of health outcomes, but not always consistently or negatively.^27–31^

### Unanswered questions and further research

Further descriptive work is required on the contribution of different causes of death, age-specific components and inequalities to the trends in Scotland. Work to understand the theoretical interaction of different hypothesised causes, and to test these theories is urgently required. In the meantime, governments at all levels should seek to provide public services according to need and sufficient social protection for all of their populations as key determinants of health. Providing effective vaccination programmes against influenza and sufficient health and social care capacity to deal with surges in demand is also required.

## Conclusion

Between 2012 and 2016 the rate of improvement in mortality markedly slowed across many high-income countries, and particularly in England & Wales, the USA, Scotland, Iceland and Northern Ireland. For this period in Scotland, the increases were only 2.5 weeks/year for women and 4.5 weeks for men. The timing of the change in mortality trend in Scotland for all ages is best estimated for men in the year to 2012 Q4 and for women in the year to 2014 Q2. Further research is required to test the range of theories for the causes of these trends, but in the meantime, governments should take action to ensure effective public services, adequate incomes, health and social care services and influenza vaccination programmes are in place.

## Competing interests

The authors declare that they have no competing interests. No funding was received for this work.

## Funding

This research received no specific grant from any funding agency in the public, commercial or not-for-profit sectors. GM, LF, JM, GW and CF are salaried by the NHS, and JR and MK are salaried by NRS.

## Contributor statement

GM drafted the manuscript. LF and JM undertook the analyses. JR and MK provided data for the segmented regression analysis. All authors made substantial contributions to editing the manuscript and approved the final draft.

---

**Appendix figure 1:**
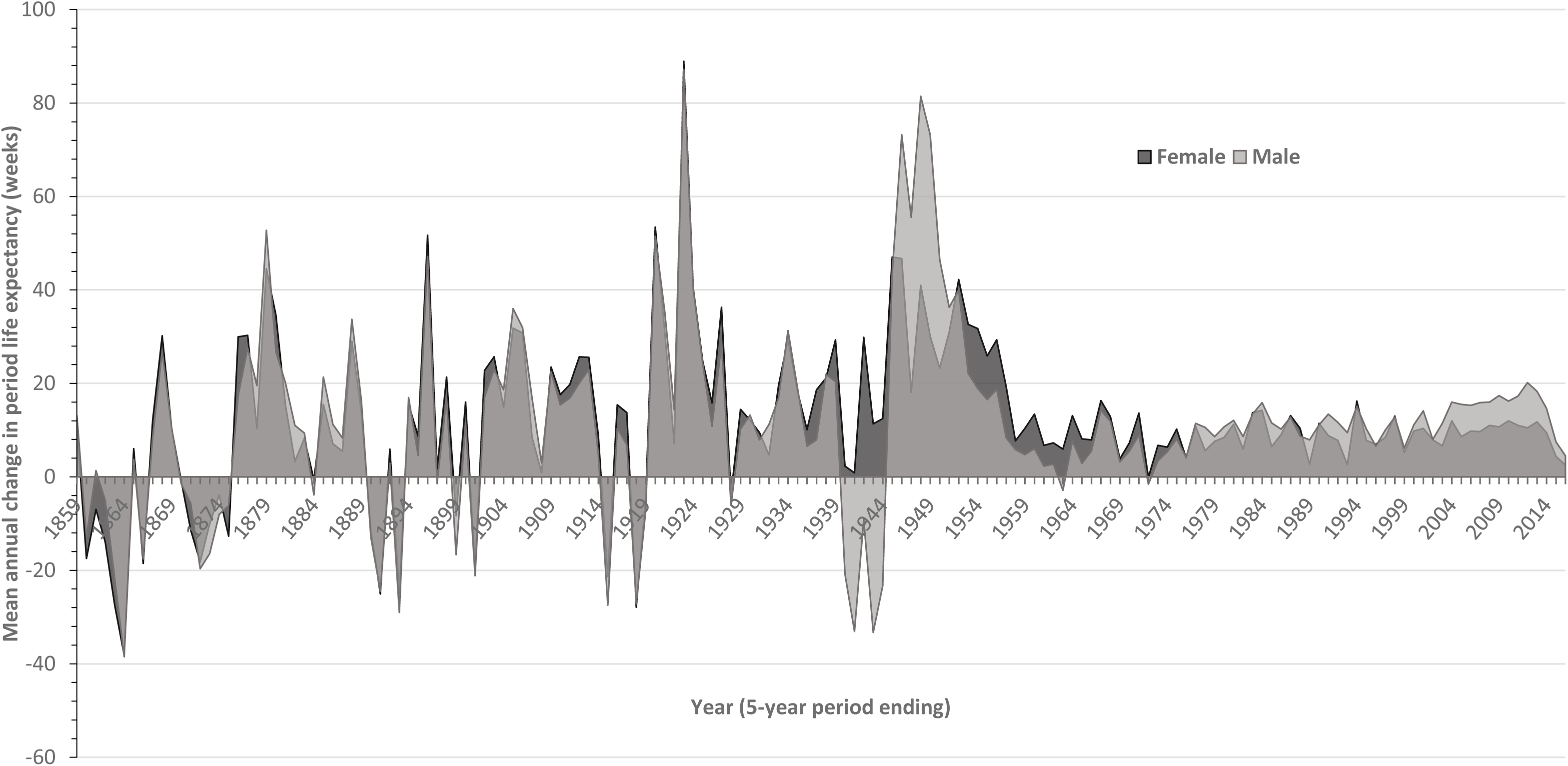
Mean annual average change in life expectancy (civilian population) for 5-year rolling periods, Scotland, males and females, 1859-2016. Data source: Human Mortality Database.

**Appendix figure 2:**
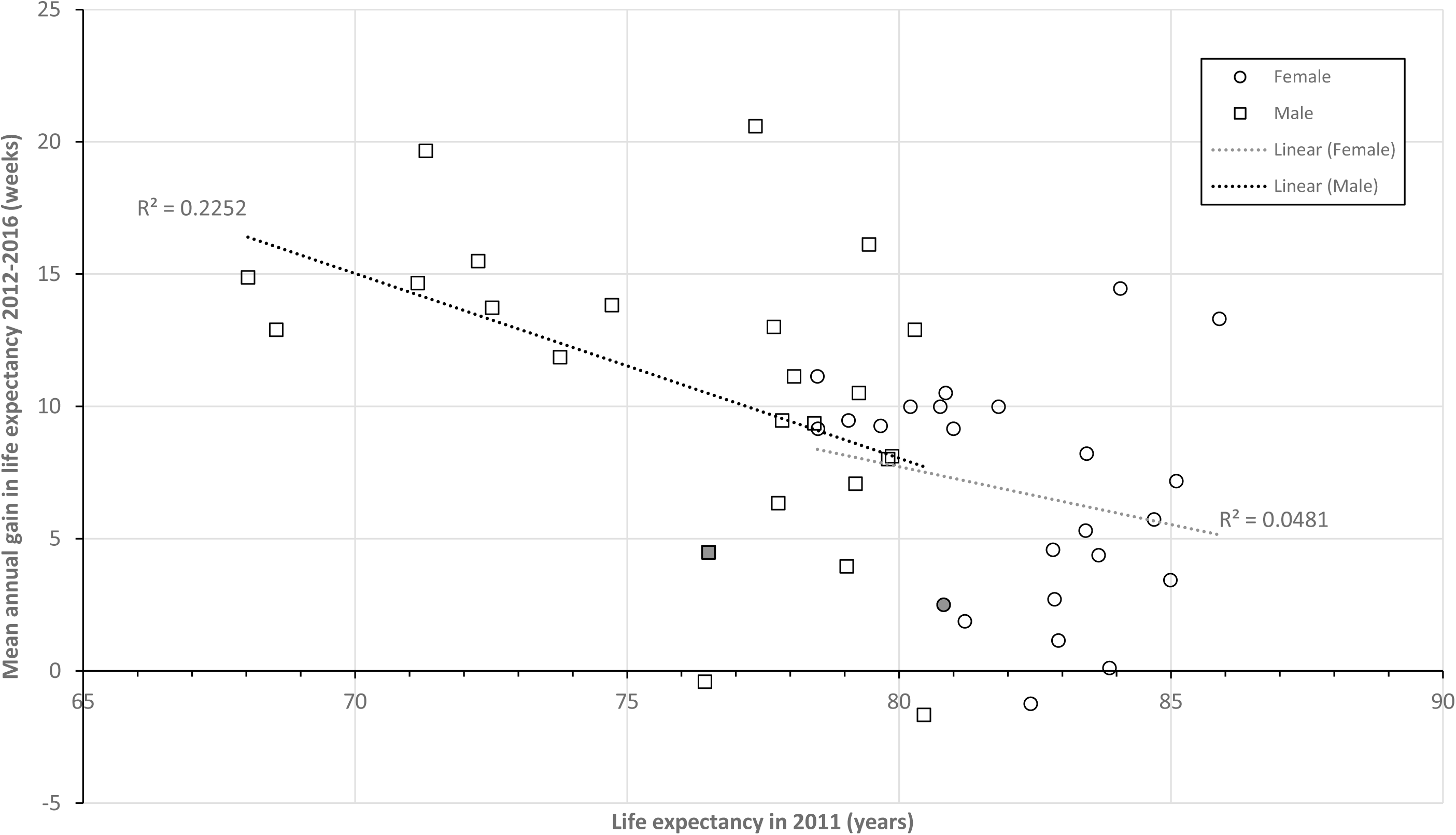
Relationship between life expectancy in 2011, and mean annual gain in life expectancy 2012-2016, for 24 high-income countries: Austria, Croatia, Czech Republic, Denmark, England & Wales, Estonia, France, Germany, Hungary, Iceland, Israel, Japan, Korea, Latvia, Lithuania, Netherlands, Northern Ireland, Poland, Scotland (indicated by shaded markers), Slovakia, Spain, Sweden, Switzerland, USA.

## Footnotes

1 24 of 42 HMD countries included. Excluded countries: No data for 2016: Australia, Belgium, Canada, Chile, Finland, Greece, Ireland, Italy, Luxembourg, New Zealand, Norway, Portugal, Slovenia, Taiwan, and Ukraine. Not high-income: Belarus, Bulgaria, and Russia.

